# Feedback Regulation between Initiation and Maturation Networks Orchestrates the Chromatin Dynamics of Epidermal Lineage Commitment

**DOI:** 10.1101/349308

**Authors:** Lingjie Li, Yong Wang, Jessica L. Torkelson, Gautam Shankar, Jillian M. Pattison, Hanson H. Zhen, Zhana Duren, Fengqin Fang, Sandra P. Melo, Samantha N. Piekos, Jiang Li, Eric J. Liaw, Lang Chen, Rui Li, Marius Wernig, Wing H. Wong, Howard Y. Chang, Anthony E. Oro

## Abstract

Tissue development results from lineage-specific transcription factors (TF) programming a dynamic chromatin landscape through progressive cell fate transitions. Here, we interrogate the epigenomic landscape during epidermal differentiation and create an inference network that ranks the coordinate effects of TF-accessible regulatory element-target gene expression triplets on lineage commitment. We discover two critical transition periods: surface ectoderm initiation and keratinocyte maturation, and identify TFAP2C and p63 as lineage initiation and maturation factors, respectively. Surprisingly, we find that TFAP2C, and not p63, is sufficient to initiate surface ectoderm differentiation, with TFAP2C-initiated progenitor cells capable of maturing into functional keratinocytes. Mechanistically, TFAP2C primes the surface ectoderm chromatin landscape and induces p63 expression and binding sites, thus allowing maturation factor p63 to positively auto-regulate its expression and close a subset of the TFAP2C-initiated early program. Our work provides a general framework to infer TF networks controlling chromatin transitions that will facilitate future regenerative medicine advances.

## INTRODUCTION

Somatic tissue development, where pluripotent stem cells progressively commit into more specialized cell types, involves dynamic changes in gene expression and chromatin organization. Cells from different lineages possess specific chromatin accessibility patterns and cis-regulatory elements (RE) that instruct lineage-specific transcription factors (TF) to precisely control their target genes (TG). While studies of individual TFs have elucidated discrete functions, detailed information is lacking about TF functions within a larger interconnected TF network. In addition, while lineage commitment requires an epigenetic transition from progenitor to terminally-differentiated cells, a paucity of information exists how stage-specific transcription factor networks interconnect to drive chromatin landscape maturation to the final committed state.

Stratified epidermal development is an ideal model system to investigate chromatin dynamic mechanisms. The epidermis represents a late ectoderm derivative, forming from lateral surface ectoderm initially specified by gradient morphogen induction by bone morphogenetic protein (BMP) and retinoic acid (RA) (Li et al., 2013; Metallo et al., 2008). Surface ectoderm is a single-layered epithelium expressing keratin 8 (K8), and keratin 18 (K18). In the presence of insulin, FGF, and EGF, surface ectoderm commits to form stratified epidermal progenitors called basal keratinocytes expressing keratin 14 (K14) and keratin 5 (K5) that are capable of producing multi-layered skin (Koster and Roop, 2007). Previous efforts have begun to identify key TFs regulating skin differentiation. The p53 family member p63 regulates keratinocyte proliferation and epidermal stratification, and loss of p63 causes skin and limb hypoplasia (Mills et al., 1999; Yang et al., 1999). However, while the role of p63 during epidermal commitment is clear, how p63 connects with upstream transcription networks that drive surface ectoderm initiation and how it ensures forward differentiation and commitment remains unclear.

An important advance in understanding epidermal TF networks comes from the ability to drive pluripotent stem cells (PSCs), including embryonic stem cells (ESCs) or induced pluripotent stem cells (iPSCs) into keratinocytes (Guenou et al., 2009; Metallo et al., 2008), thus enabling the collection of genome-wide regulatory information from cells at corresponding stages. Recently, we and others have used stem cell technologies to successfully generate patient-specific, genetically-corrected iPSC-derived graftable keratinocyte sheet for treatment of epidermolysis bullosa, a genetic blistering disease caused by *Collagen VII* mutation (Sebastiano et al., 2014; Umegaki-Arao et al., 2014; Wenzel et al., 2014). While these findings provide hope for tissue replacement therapies, a major roadblock remains the understanding and improvement of the differentiation process that will increase its efficiency and specificity to a level compatible with clinical manufacturing. Toward this end, dissecting the genome-wide regulatory landscape during differentiation remains critical for understanding lineage commitment in epidermal development.

Here, we use a defined feeder-free, xeno-free ES differentiation system and a newly-described network inference modeling algorithm to identify the interconnecting TF networks during two major epigenetic transition periods. Subsequent functional studies uncover the surprising finding that a single factor, TFAP2C, drives skin differentiation by initiating the surface ectoderm chromatin landscape and inducing the maturation factor p63; p63, in turn, matures the chromatin landscape into stratified epithelium and inhibits select aspects of the TFAP2C surface ectoderm network. Our work defines the regulatory landscape during human epidermal lineage commitment and elucidates key regulatory principles that enable somatic tissue development and future stem cell-based regenerative therapy.

## RESULTS

### Epigenomic Profiling Identifies Key Transitions during Epidermal Commitment

We capitalized on our defined in vitro differentiation protocol (Sebastiano et al., 2014) to detail the transcription and chromatin dynamics at each stage of differentiation. In brief, human pluripotent ESCs were treated with RA and BMP4 to induce surface ectoderm differentiation for seven days, producing a simple epithelial cell (K8^+^/K18^+^). The surface ectoderm progenitor cells were then treated with defined keratinocyte serum-free medium (DKSFM, containing insulin, EGF and FGF) to drive epidermal lineage maturation, consisting of cell death, migration, and epithelial colony formation of pure hESC-derived basal keratinocytes (H9KC) (Figures 1A and S1A). H9KCs possessed similar morphology and marker gene expression as somatic foreskin keratinocytes (K14^+^p63^+^K18^-^) (Figures 1B and S1A) and formed stratified epidermal layers in organotypic cultures (K10^+^ and Loricrin^+^) (Figure 1C), providing us with a useful model for skin differentiation.

**Figure 1.**
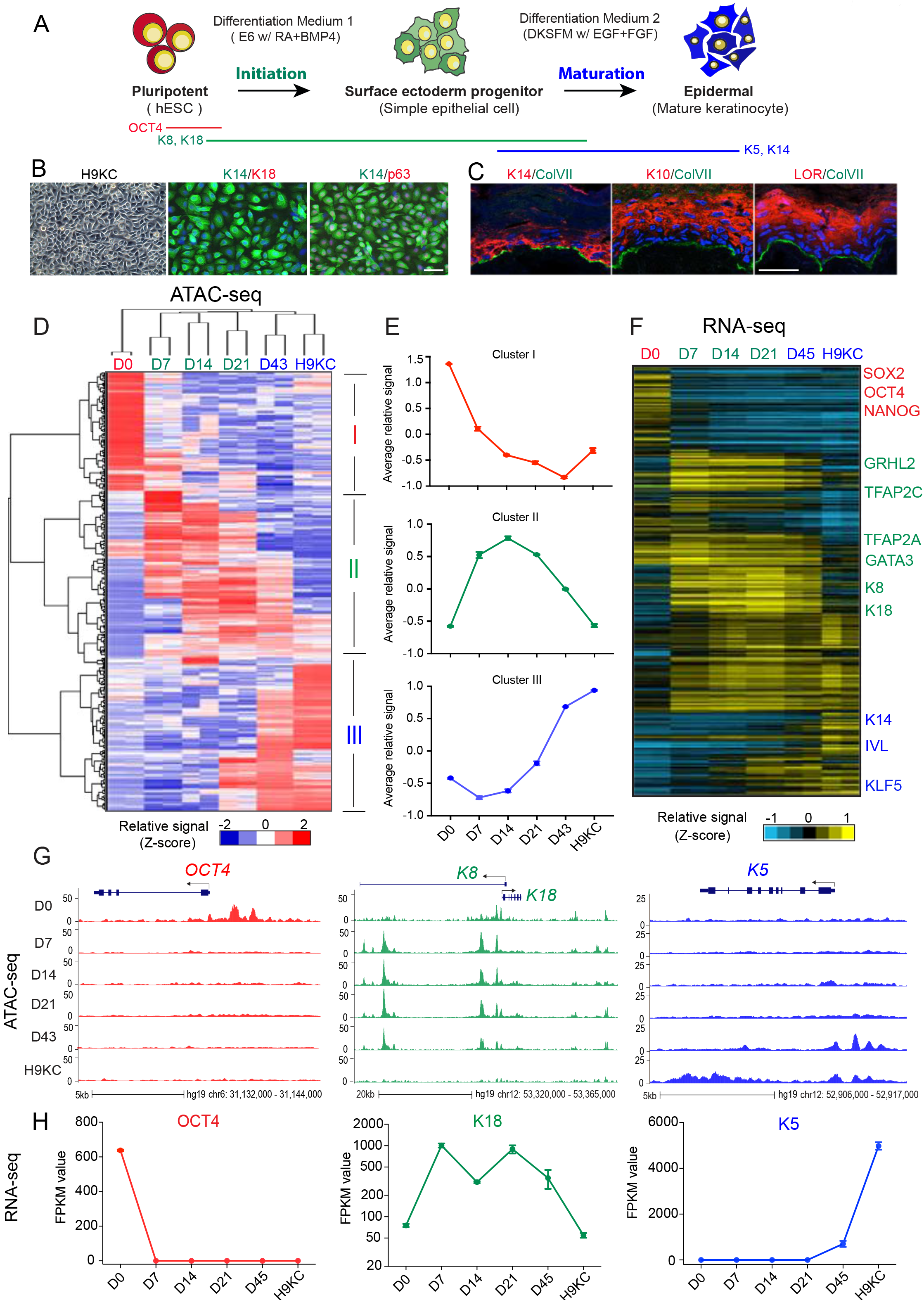
Accessible Chromatin and Transcriptome Landscapes Identify Three Major Chromatin Stages during Epidermal Lineage Commitment. (A) Schematic overview of epidermal differentiation from human embryonic stem cells (hESC). (B) Light image and immunofluorescence (IF) staining of keratinocytes derived from H9 hESCs (H9-KC). The cells were stained with keratinocyte-specific markers K14 (green), p63 (red), and simple epithelial marker K18 (red). The nuclei were stained with DAPI (blue). Scale bar: 50 μm. (C) Reconstruction of stratified epidermis with hESC-derived keratinocytes in organotypic culture. IF staining showing the maker localization in the H9KC organotypic epidermis. Basal epidermal marker: K14 (red); suprabasal epidermal marker: K10 (KRT10, red), LOR (Loricrin, red); basement membrane marker: ColVII (Collagen VII, green). The nuclei were stained with DAPI (blue). Scale bar: 25μm. (D) Heat map of differential open chromatin regulatory elements characterized from ATAC-seq at each time point during epidermal differentiation. Each column is a sample and each row is a regulatory element. Hierarchical clustering of both sample and element yields three big clusters of element (labeled as cluster I, II, and III) and three major groups of sample (group 1: D0, red; group 2: D7, D14, D21, green; and group 3: D43, H9KC, blue). The color bar shows the relative ATAC-seq signal (Z-score of normalized read counts) as indicated. (E) The trend of signal changes of the three clusters identified from ATAC-seq in (D). (F) Heat map of the expression changes of the genes containing differential ATAC-seq signals at their promoters. The color bar shows the relative expression value [Z-score of FPKM (Fragments Per Kilobase of transcript per Million mapped reads)] from the RNA-seq. (G) Normalized ATAC-seq profiles at *OCT4*, *K8-K18* and *K5* loci, representing the dynamic ATAC-seq signal changes of the three clusters identified in (D) and (E) respectively. (H) Gene expression changes of OCT4, K18, and K5. The FPKM values of each gene at all the time points are shown. See also **Figure S1.**

We characterized the epigenetic changes occurring over the 60-day skin differentiation by identifying open chromatin regulatory elements and associated gene expression changes at 6 specified time points using Assay for Transposase Accessible Chromatin with high-throughput sequencing (ATAC-seq) (Buenrostro et al., 2013) and RNA-seq methods (Figure S1B). We identified a total of 336,268 open chromatin REs across all samples, and selected 188,850 that were differentially enriched at specific stages (Figures 1D and S1C). Similarly, we characterized the total and stage-specific transcriptome and focused on 9,362 transcripts that defined each time point. We used the StepMiner algorithm with D0 and H9KC data to determine whether differentiation occurred continuously or in a stepwise fashion (Sahoo et al., 2007). We observed progressive reduction of pluripotent-specific sites and acquisition of keratinocyte-specific sites over time (Figures S1E-F). The associated-gene expression pattern follows a similar trend (Figures S1E-F), suggesting a gradual, rather than stepwise, maturation to keratinocytes.

Hierarchical clustering of the differential peaks led us to define three major sample groups that describe the changing epigenetic landscape: group 1 (early stage: D0), group 2 (middle stage: D7, D14 and D21) and group 3 (late stage: D43 and H9KC) (Figure 1D). Peak clusters correspond with sample groups that characterize the chromatin accessibility features of pluripotent cell, surface ectoderm progenitor and mature keratinocyte with the terms “initiation” and “maturation” used to describe the sequential transitions of each states (Figure 1A). Cluster I contains elements with higher chromatin accessibility signals in pluripotent cells that close upon differentiation (Figures 1D-F, and S1D). As a typical gene locus, both the chromatin accessibility and mRNA abundance of the pluripotent factor OCT4 (POU5F1) decrease from D0-D7 (Figures 1G-H), reflecting a transition from pluripotent state to non-neural ectoderm differentiation. Cluster II reflects the emergence of chromatin accessibility of genes associated with surface ectoderm initiation from D7-21 (Figures 1D-F, and S1D). This cluster remains open until keratinocyte maturation, at which point a majority of the loci close. K8, K18 are typical markers for simple, but not stratified epithelial cells and the accessible signals at *K8-K18* locus and the mRNA expression of K18 are specifically higher in the middle stage (Figures 1G-H). Finally, Cluster III contains elements with increased accessibility changes and higher associated gene expression during late stage of differentiation as definitive keratinocytes emerge (D43 and H9KC) (Figures 1D-F, and S1D). A typical example is the keratinocyte-specific marker K5 whose locus opens and gene expression increase from stage D43 to H9C (Figures 1G-H). We conclude that addition of RA/BMP drives ES cells through 3 major stages of differentiation defined by transitions of Initiation and Maturation.

### TF-Chromatin Transcriptional Regulatory Networks during Epidermal Commitment

To create a comprehensive TF network of keratinocyte differentiation, we took advantage of the observation that accessible chromatin regions represent unique cell state markers that are recognized by specific TFs and developed an algorithm that linked changes in accessible chromatin using ATAC-seq, the predicted REs for specific TFs, and the TF activity associated with a transcription start site as revealed through RNA sequencing. First, we created a master list of all TF binding motifs in the changing accessible regions at each time point by HOMER software, identifying a total of 39 motifs from all stages. Next through filtering by motif enrichment score and TF expression changes, we identified 17 key TF motifs most correlated with the dynamic chromatin accessibility landscape (Figure 2A and Table S1).

To provide a detailed view of how TFs and open REs work together to dynamically affect gene expression, we employed our recently described network inference framework that combines gene expression and chromatin accessibility data with accessible TF binding motifs (Duren et al., 2017). A major hurdle for network modeling has been the discovery that most of the chromatin changes seen during keratinocyte differentiation occur in the distal regulatory regions (enhancers) rather than the promoters, an observation we confirm in our own keratinocyte differentiation analysis (Figure S1C), making it difficult to associate RE to TG and assign transcriptional start sites to local chromatin changes. To overcome this challenge, we incorporated publicly available chromosome conformation data and co-occurrence information of distal DNase I hypersensitive sites (DHSs) and promoter DHSs across 79 diverse cell types from ENCODE (Rao et al., 2014; Thurman et al., 2012), which significantly enhanced the accuracy of gene assignment in our model (Figures 2B, S2B and Table S2). We focused on the two transition periods, the Initiation phase (i.e. the transition from Stage 1 to 2) and Maturation phase (the transition from Stage 2 to 3). Each transition period represents changes in chromatin accessibility associated with a TF-associated RE and a particular TG. We assumed that the rate of transcription of a TG depends on TFs bound to regulatory elements that are open at that time point (Figure 2B). This TF-RE-TG triplet simplification allowed us to integrate all investigated genomic features. We then quantitatively ranked all the TF-RE-TG triplets and selected the high confident connection triplets to generate a network for representation (Figures 2B, S2A, S2B and Table S3). In the early differentiation stage (“Initiation”) where hESCs lose the pluripotency and differentiate into committed surface ectodermal cells, AP2 factors (TFAP2A, TFAP2C), GATA3 and GRHL2 gain chromatin accessibility of TF binding sites and transcriptional activity, while pluripotent factors POU5F1, NANOG, and SOX2 lose chromatin accessibility of TF binding sites and TG expression (Figures 2C and S2C). By contrast, keratinocyte maturation correlates with the increased accessibility of p63, KLF4 and AP1 binding sites and associated gene expression, while displaying decreased accessibility for key surface ectoderm regulators like TFAP2C and GATA3 (Figures 2C and S2C). To validate the network finding, we investigated individual TF occupancy and TG expression by ChIP-seq and RNA-seq, and the results show the correlated efforts of initiation and maturation TF networks (Figure S2D). We conclude that inference network modeling provides a ranked TF network that describes key transitions during keratinocyte differentiation.

**Figure 2.**
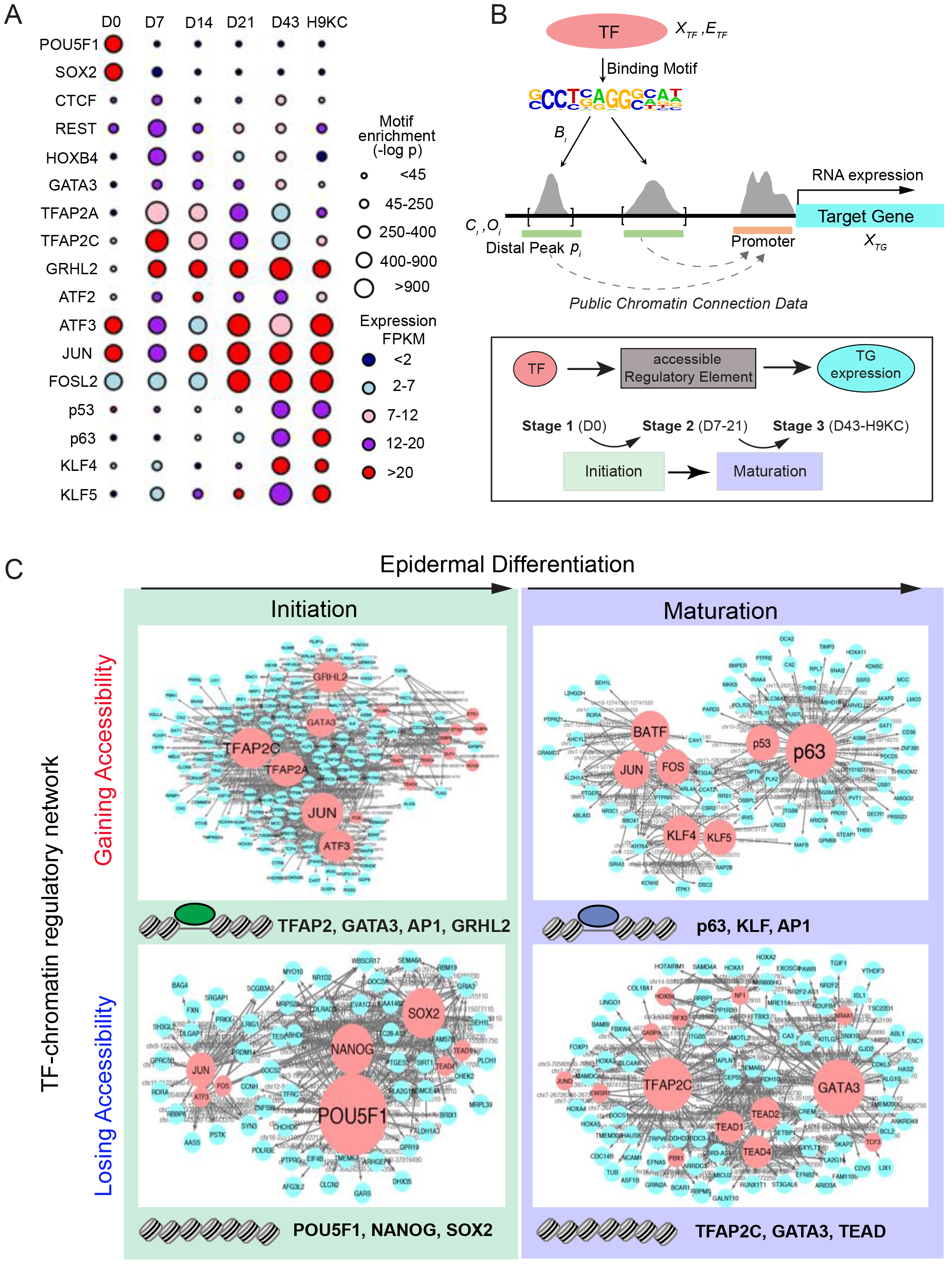
Identification of Master TFs Driving Surface Ectoderm Initiation and Keratinocyte Maturation by TF-Chromatin Transcriptional Regulatory Network. (A) TF motifs identified from differential ATAC-seq peaks at each time point. The circle size represents different level of motif enrichment (measure by −log_10_ p-value); and the color represents the expression level (FPKM value) of each TF in RNA-seq. (B) Schematic overview of the method of constructing the TF-chromatin transcriptional regulatory network. Top panel: establishment of a connection between TF and target gene (TG) through open chromatin binding. The ATAC-seq peak regions were scanned to find specific TF binding motifs located in the distal or promoter regions of a gene; public available chromatin connection data was used to annotate the possible connection between distal peak to promoter-associated peak (dashed line with arrow). Bottom panel: The triple elements, i.e. TF-accessible regulatory elements (RE, i.e. ATAC-seq peaks)-TG expression, were ranked by the coherence among genomic features and extracted through a statistical model to build the regulatory network. We investigated and ranked their feature changes (Figure S2A and S2B) during two major transition events, Initiation and Maturation; which were identified by clustering of ATAC-seq samples during differentiation. (C) TF-chromatin transcriptional regulatory networks identified from the chromatin regions gaining or losing accessibility in the Initiation and Maturation process. In the network, the red and blue nodes represent TF and TG; and the gray edge represents the accessible regulatory element which was bound by TF to regulate TG expression. Larger size of TF nodes represent more TG connections. Top ranked TFs were listed at the bottom of each network. Note: TF family name was used to represent several members. See also **Figure S2.**

### TFAP2C Alters Chromatin Landscape to Drive Surface Ectoderm Differentiation

Using the top regulators identified by the network, we performed functional genomics studies to dissect the hierarchical relationships of TFs during surface ectoderm initiation and keratinocyte maturation. We established doxycycline (Dox)-inducible piggy bac hESC lines of the major TFs to investigate ectopic expression, and CRISPR-treated lines to assay loss-of-function phenotypes with the goal of determining which TF could replace the addition of RA/BMP. Surprisingly, overexpression of TFAP2C (“TetO-TFAP2C”, Figure 3A) which has the highest fold change expression and ranking position in the early network (Figures 2A and 2C), induced a flattened and adhesive epithelial-like morphology (Figure S3A) in the absence of RA/BMP. Compared to control cells, TFAP2C-overexpressing cells induced elevated K18 protein within filamentous cytoplasmic structures, a typical simple epithelial staining pattern seen in RA/BMP4 induced surface ectodermal cells (Figure 3B). While sufficient to drive surface ectoderm differentiation, we determined whether TFAP2C is also necessary. We attempted to remove TFAP2C in ESCs by CRISPR/Cas9, but failed to generate homozygous knockout after several rounds of gene targeting, discovering a growth defect in ESCs upon siRNA-mediated TFAP2C gene expression knock-down (Figures S3B-C). Nevertheless, in the heterozygous knockout cell line (TFAP2C ^+/−^), TFAP2C protein levels are 20% of wildtype at D7 and K18 protein level fail to increase (Figure 3B). In addition, the mRNA level of many surface ectodermal genes decline upon TFAP2C loss (Figure 3C), supporting an essential role in early commitment. Taken together, we conclude that TFAP2C is necessary and sufficient to induce the surface ectodermal phenotype.

As TFAP2C-induced surface ectoderm differentiation mimicked early RA/BMP-induced differentiation, we compared in depth the expression and chromatin accessibility changes upon TFAP2C activation. RNA-seq and qRT-PCR analysis indicated that expression levels of most surface ectodermal makers, such as K8, TFAP2A, GATA3, and GRHL2, were all increased significantly (Figure 3D) with a concomitant decrease in the expression of pluripotency or other germ layer markers (Figure 3E). Consistent with this analysis, gene set enrichment analysis (GSEA) also confirms a significant enrichment (NES=1.42, p<0.001) of surface ectoderm genes (Qu et al., 2016)(Figure S3D), and gene expression changes display high correlation coefficients (R=0.8967, p<0.0001) compared to D7 cells (Figure 3F).

Chromatin accessibility changes also support TFAP2C-inducible surface ectoderm commitment. First, principle component analysis (PCA) comparing the similarity of ATAC-seq signals from TFAP2C over-expressed cells (hereafter “TetO-TFAP2C-D7 Dox+”) at Day 7 with the different stages of RA/BMP differentiation showed that TFAP2C-D7 Dox+ cells match those from D7, but are distinct from D0 or later time points (D43, and H9KC) (Figure 3H). Next, direct comparison of the open chromatin signatures of TetO-TFAP2C-D7 Dox+ and D7 cells revealed a high correlation (R=0.7714, p<0.0001) (Figure 3G). GO enrichment terms for genes associated with gained accessibility were enriched in adherens junction, tight junction, hippo signaling, desmosome assembly, while genes associated with lost accessibility were enriched in neuron differentiation, pattern specification process, etc. (Figures 3I and S3F). Detailed analysis shows increased ATAC-seq signals around the transcription start site (TSS) in surface ectodermal genes after TFAP2C overexpression (Figure S3E), indicating the activation of gene transcription at loci such as *K8-K18* and the Initiation network TF *GATA3* (Figures 3E and 3J).

**Figure 3.**
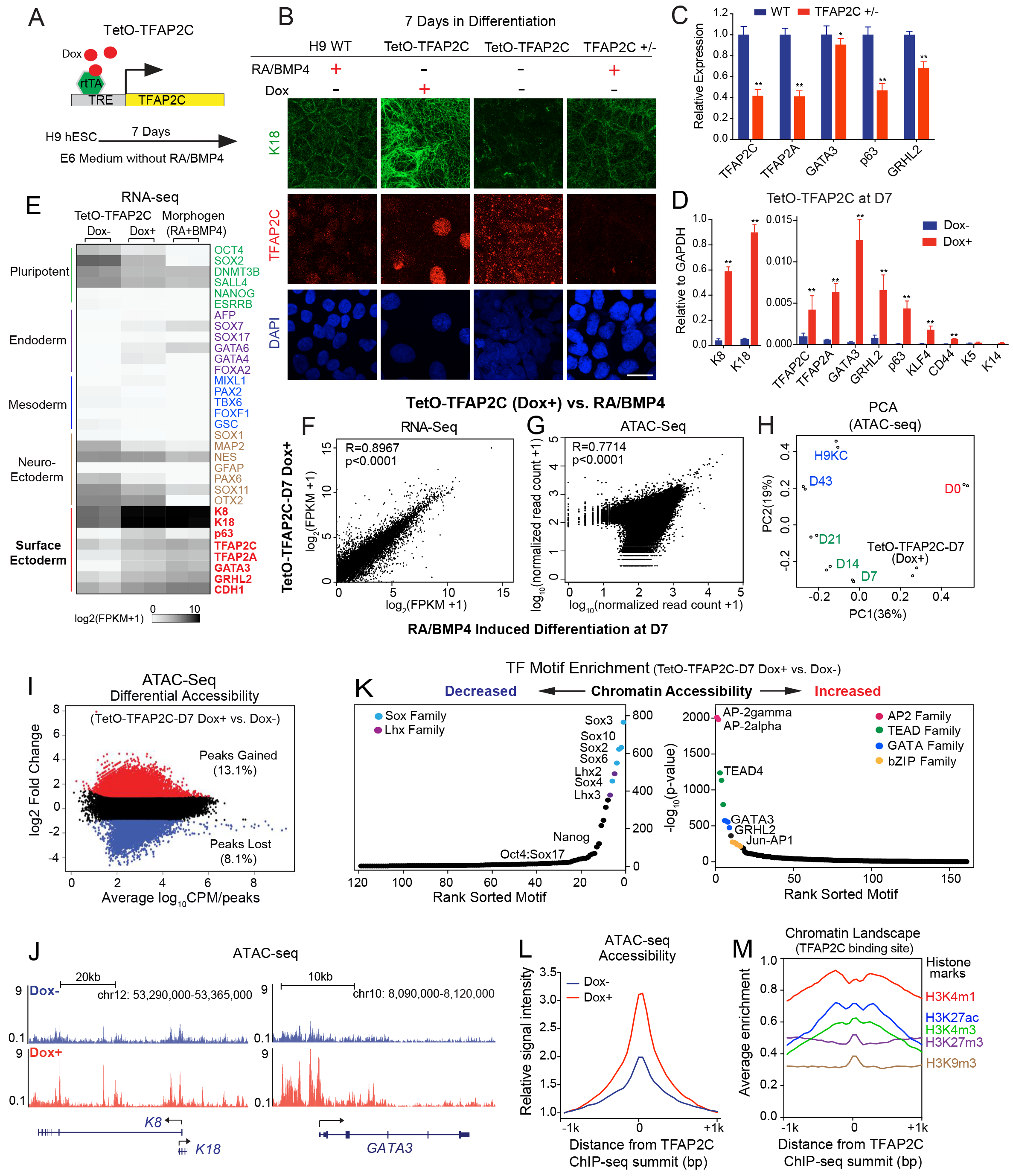
TFAP2C Initiates the Chromatin Landscape to Induce Surface Ectoderm Differentiation. (A) Schematic representation of the TetO-TFAP2C inducible expression system in H9 hESC. A cassette containing the TFAP2C cDNA is under the control of the tetracycline − responsive element (TRE). Upon Doxycycline (Dox) induction, the expression of TFAP2C is activated by reverse tetracycline transactivator rtTA. TetO-TFAP2C cells were treated with Dox in E6 medium without adding morphogen RA/BMP4 and the differentiation effect was evaluated in the following assays. Same cells without Dox induction served as the control. (B) IF staining of K18 and TFAP2C in TetO-TFAP2C cells with and without Dox induction, and cells treated with RA/BMP4 from H9 WT and TFAP2C heterozygous (+/−) hESCs. Scale bar: 20μm. (C) Gene expression changes in early differentiation upon TFAP2C loss of function. Wild type (WT) and CRISPR/Cas9 mediated heterozygous knockout of TFAP2C (TFAP2C+/−) hESCs were induced into surface ectoderm progenitor cells by RA/BMP4 for 7 days. qRT-PCR values were normalized to the values in WT group. Mean ± S.D. is shown; significant difference with *p<0.05, **p<0.01 relative to WT determined by t-test (n=3). (D) Gene expression changes in TetO-TFAP2C cells upon Dox induction. The cells were cultured in E6 basal medium with or without Dox for 7 days for comparison (Dox+ vs. Dox-). qRT-PCR values were normalized to the values of internal control GAPDH. Mean ± S.D. is shown; significant difference with **p<0.01 relative to control (Dox-) determined by t-test (n=3). (E) Heat map of gene expression changes from the RNA-seq in “TetO-TFAP2C” cells with/without Dox addition, and differentiated cells induced by morphogens (RA+BMP4) for 7 days. Germ layer specific genes, including pluripotent, endoderm, mesoderm, neuro-ectoderm and surface ectoderm, were examined. All the surface ectodermal genes have increased expression upon TFAP2C over expression (Dox+) and morphogen induction compared to that in the control cells (Dox-), while there are decreased or non-significant changes in other germ layer genes. (F) Scatter plot of gene expression from RNA-seq [expressed as log_2_(FPKM+1)] in early differentiated cells by TFAP2C activation (TetO-TFAP2C Dox+, y axis), versus by RA/BMP4 induction (x, axis) at day7 with pearson correlation (R) value displayed (R=0.8967, p<0.0001). (G) Scatter plot of ATAC-seq counts from combined peaks in early differentiated cells by TFAP2C activation (TetO-TFAP2C Dox+, y axis), versus by RA/BMP4 induction (x, axis) at day7 with pearson correlation (R) value displayed (R=0.7714, p<0.0001). (H) Principle component analysis (PCA) of the ATAC-seq samples from early differentiated cells with TFAP2C activation [TetO-TFAP2C-D7 (dox+)], and samples at all time points during normal differentiation. (I) Scatter plot of differential accessibility between TetO-TFAP2C-D7 Dox+ and Dox-cells. y-axis: log_2_ fold change in reads per accessible region (Dox+ vs. Dox-); y-axis: average log_10_ count per million reads (CPM) per accessible region (i.e. peak) in ATAC-seq. The values of log_2_ fold change >1, or < −1 are labeled as “peak gained” or “peak lost” with respective percentage and color. (J) Genome browser tracks of normalized ATAC-seq signal showing increased chromatin accessibility at *K8-K18* and *GATA3* loci upon TFAP2C-induced early differentiation (Dox+ vs. Dox-). (K) Enrichment of TF motifs in the regions with decreased (left) or increased (right) chromatin accessibility in TetO-TFAP2C-D7 Dox+ cells comparing with Dox-cells. The y-axis is −log_10_ p-value of a motif enrichment score, which is sorted from largest to smallest. The x-axis is the ranking number of sorted motif. Motifs belong to one TF family are labeled with same color and are indicated in the top corner of each panel. (L) Average enrichment of ATAC-seq chromatin accessibility within −1/+1kb from TFAP2C binding site in cells with or without Dox induction. (M) Average enrichment of individual histone marks within −1/+1kb from TFAP2C binding sites in TFAP2C over-expressed cells. See also **Figure S3.**

Importantly, TFAP2C not only induces the expression of TFs in the Initiation network but also activates the binding sites for them. TF motif enrichment with the differential accessible regions shows the most highly enriched motifs were the binding sites of AP-2 family (AP-2gamma, AP-2alpha), followed by TEAD family (TEAD4, TEAD1), GATA family (GATA3, GATA2), GRHL2, and bZIP family (JUN) (Figure 3K and S3G), while the most closed are sites for TFs responsible for ESC maintenance or other lineage specification, such as SOX family (Sox3, Sox2), Lhx family (Lhx2, Lhx3), and Nanog (Figure 3K). Subsequent foot printing analysis using protein interaction quantification (PIQ) software calculated the probability of occupancy for each candidate binding site in the genome (Sherwood et al., 2014), and revealed a statistically significant increase in the binding of such Initiation network TFs upon TFAP2C overexpression at D7 (Figure S3H). Therefore, TFAP2C is able to activate the expression and binding activity of the early network TFs, thus activating the initiation network in surface ectoderm differentiation.

TFAP2C action could be direct, or indirect through inducing expression of Initiation network TFs that could in turn open chromatin binding sites. To distinguish between these possibilities, we performed TFAP2C ChIP-seq analysis in D7 TFAP2C over-expressed cells and identified ~10,000 TFAP2C binding sites that demonstrate high enrichment for the AP-2alpha and AP-2gamma motifs, and are predominantly located in intergenic and intron regions at a distance from the gene TSS (Figures S3I-K). Consistent with a direct role, we found significantly increased chromatin accessibility around TFAP2C binding sites upon TFAP2C over-expression (Figure 3L). In addition, TFAP2C-bound regions have a higher enrichment of active histone marks such as H3K4m1, H3K27ac and H3K4m3 (Figure 3M). Collectively, these results support the conclusion that TFAP2C binds to the active regulatory regions to increase genome-wide chromatin accessibility changes associated with surface ectoderm lineage commitment.

### TFAP2C-Induced Surface Ectoderm Cells Produce Functional Keratinocytes

Since TFAP2C induces surface ectodermal differentiation by activating the endogenous early TF-chromatin transcriptional regulatory network, we determined whether the committed progenitor cells had the potential to develop into mature keratinocytes. We treated TFAP2C-induced progenitor cells (hereafter “TetO-TFAP2C-D7”) with keratinocyte maturation medium and withdrew Dox after 10 days, to investigate the long-term differentiation ability (Figure 4A). Intriguingly, we found that TetO-TFAP2C-D7 possessed a 5-fold higher yield of keratinocyte colonies (hereafter “TetO-TFAP2C-KC”) than RA/BMP induced keratinocytes, colonies whose cells possessed typical cobblestone morphology, K14, ITGA6 and p63 staining, and lacked simple epithelial marker K18 (Figure 4C). Similarly, qRT-PCR analysis also confirmed elevated transcription of mature keratinocyte, but not simple epithelial makers, and decreased expression of the Initiation network TFs TFAP2C, GATA3 and GRHL2 (Figure 4B). In addition, fluorescence activated cell sorting (FACS) (Figure S4A) shows that majority of “TetO-TFAP2C-KC” cells are K14^+^K18^-^ (95%), a pattern similar to that in normal human keratinocyte (NHK, K14^+^K18^-^, 95%), indicating a near complete transition from simple epithelial cells to mature keratinocytes.

Further analysis validated the ability to form a stratified epithelium. Foreskin and RA/BMP induced keratinocytes respond to high calcium stimulation, which results in cellular structural changes and increased expression of markers specific for stratification process (K1, K10, IVL, FLG), and depression of basal layer markers (p63, ITGA6, ITGB4). TFAP2C-induced keratinocytes demonstrate rapid calcium-induced cornification (Figures S4B-C). Using an *in vitro* skin reconstitution assay with decellularized dermis, we found that the cells were able to form all stratified layers with positive staining of respective basal, squamous, and granular layer markers (Figure 4D), indicative of the ability to create a stratified epidermal tissue. Hierarchical clustering of ATAC-seq signals from RA/BMP-induced differentiation and TetO-TFAP2C-KC revealed that the chromatin landscape of TetO-TFAP2C-KC is close to that in late differentiated cells (D43, H9KC), and distinct from cells at earlier stages (Figure 4E). In addition, ATAC-seq accessibility (R=0.8736, p<0.0001) (Figure 4F), and RNA-seq signatures (R=0.8252, p<0.0001) (Figure 4G) between TetO-TFAP2C-KC and H9KC are highly correlated and include enrichment of motifs for Maturation network TFs, and reduced enrichment for Initiation network TFs (Figures 4H-I, and S4D). Subsequent PIQ analysis revealed a statistically significant increase in the binding of most Maturation network TFs in TetO-TFAP2C-KC (Figure S4E). Significantly, we found a decrease of ATAC-seq signal in the regulatory regions of *K8-K18* locus, and a strong increase in *K14* locus in TetO-TFAP2C-KC vs. TetO-TFAP2C-D7 (Figure 4J). We conclude that TFAP2C-induced surface ectodermal progenitors are competent to further differentiate into functional keratinocytes, which is accomplished by the transition of chromatin landscape from the Initiation to the Maturation network.

**Figure 4.**
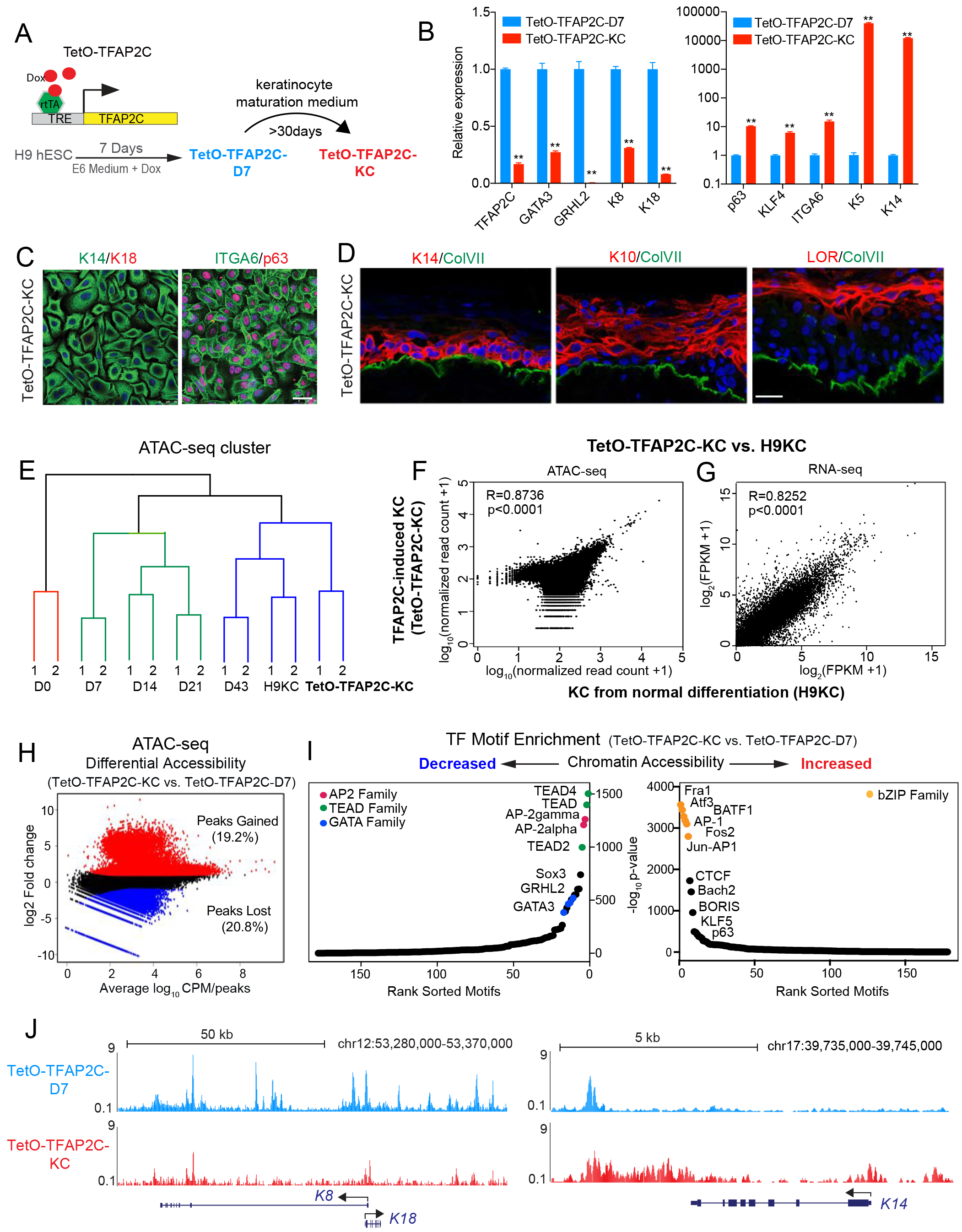
TFAP2C-Induced Surface Ectoderm Progenitors Are Competent to Produce Functional Keratinocyte in Maturation Medium. (A) Schematic diagram showing the experimental procedure to test whether TFAP2C-induced surface ectoderm progenitor cells (TetO-TFAP2C-D7) can further differentiate into functional keratinocytes (TetO-TFAP2C-KC) in maturation medium for over 30 days. (B) Gene expression changes in late differentiation by comparing TetO-TFAP2C-KC vs. TetO-TFAP2C-D7. qRT-PCR values were normalized to the values in the group of TetO-TFAP2C-D7. Mean ± S.D. is shown; significant difference with **p<0.01 relative to “TetO-TFAP2C-D7” determined by t-test (n=3). (C) IF staining of K14, K18, ITGA6 and p63 in keratinocytes derived from TFAP2C-induced differentiation (TetO-TFAP2C-KC). Nuclei were stained with DAPI (blue). Scale bar: 25μm. (D) Reconstruction of stratified epidermis with TFAP2C-induced keratinocytes in organotypic culture. IF staining of K14, K10, LOR and ColVII are shown. Nuclei were stained with DAPI (blue). Scale bar: 25μm. (E) Hierarchical clustering of “TetO-TFAP2C-KC” and samples from normal differentiation using their chromatin accessibility similarities from ATAC-seq analysis. The three major clusters are highlighted in different colors. (F) Scatter plot of ATAC-seq counts from combined peaks in TFAP2C-induced keratinocytes (TetO-TFAP2C-KC, y axis), and keratinocytes from normal differentiation (H9KC, x axis) with pearson correlation (R) value displayed (R=0.8736, p<0.0001). (G) Scatter plot of gene expression from RNA-seq [expressed as log_2_(FPKM+1)] in keratinocyte by TFAP2C induction (TetO-TFAP2C-KC, y axis), versus keratinocyte from normal differentiation (H9KC, x axis) with pearson correlation (R) value displayed (R=0.8252, p<0.0001). (H) Scatter plot of differential accessibility between “TetO-TFAP2C-KC” and “TetO-TFAP2C-D7” cells. y-axis: log_2_ fold change in reads per accessible region (TetO-TFAP2C-KC vs. TetO-TFAP2C-D7); x-axis: average log_10_ CPM per accessible region (i.e. peak) in ATAC-seq. The values of log_2_ fold change >>1, or <-1 are labeled as “peak gained” or “peak lost” with respective percentage and color. (I) Enrichment of TF motifs in the regions with decreased (left) or increased (right) chromatin accessibility in “TetO-TFAP2C-KC” cells comparing to that “TetO-TFAP2C-D7” cells. The y-axis is −log_10_ p-value of a motif enrichment score, which is sorted from largest to smallest. The x-axis is the ranking number of sorted motif. Motifs belong to one TF family are labeled with same color and are indicated in the top corner of each panel. (J) Genome browser tracks of normalized ATAC-seq signal showing decreased chromatin accessibility at *K8-K18* locus and increased accessibility at *K14* locus in TFAP2C-induced mature keratinocytes (TetO-TFAP2C-KC) comparing to that in early progenitor cells (TetO-TFAP2C-D7). See also **Figure S4.**

### TFAP2C Requires p63 to Drive Keratinocyte Maturation

As the TetO-TFAP2C-KCs showed a decrease in Initiation network TFs and an increase in the Maturation network TFs (Figure 2C, S2C, 4I and S4E), we functionally interrogated the maturation-associated TF required to mature cells into keratinocytes. p63 demonstrated the desired characteristics. We knocked out the p63 gene in TetO-TFAP2C cells by CRISPR/Cas9 and evaluated its effect on TFAP2C-induced differentiation (Figure 5A). TetO-TFAP2C+p63KO cells with Dox displayed typical epithelial morphology (D7, Figure 5B) and high gene expression of surface ectoderm markers by IF and qRT-PCR (Figures 5C-D), indicating that p63 is not necessary for TFAP2C-induced early initiation. By contrast, when exposed to epidermal maturation media, TetO-TFAP2C+p63KO cells underwent apoptosis rather than keratinocyte maturation (D50, Figure 5B), demonstrating p63-dependent survival. Moreover, mutant cells exhibited persistent surface ectoderm markers and lack of mature keratinocyte markers (Figures 5E-F), further confirming the p63-dependence of the maturation process.

**Figure 5.**
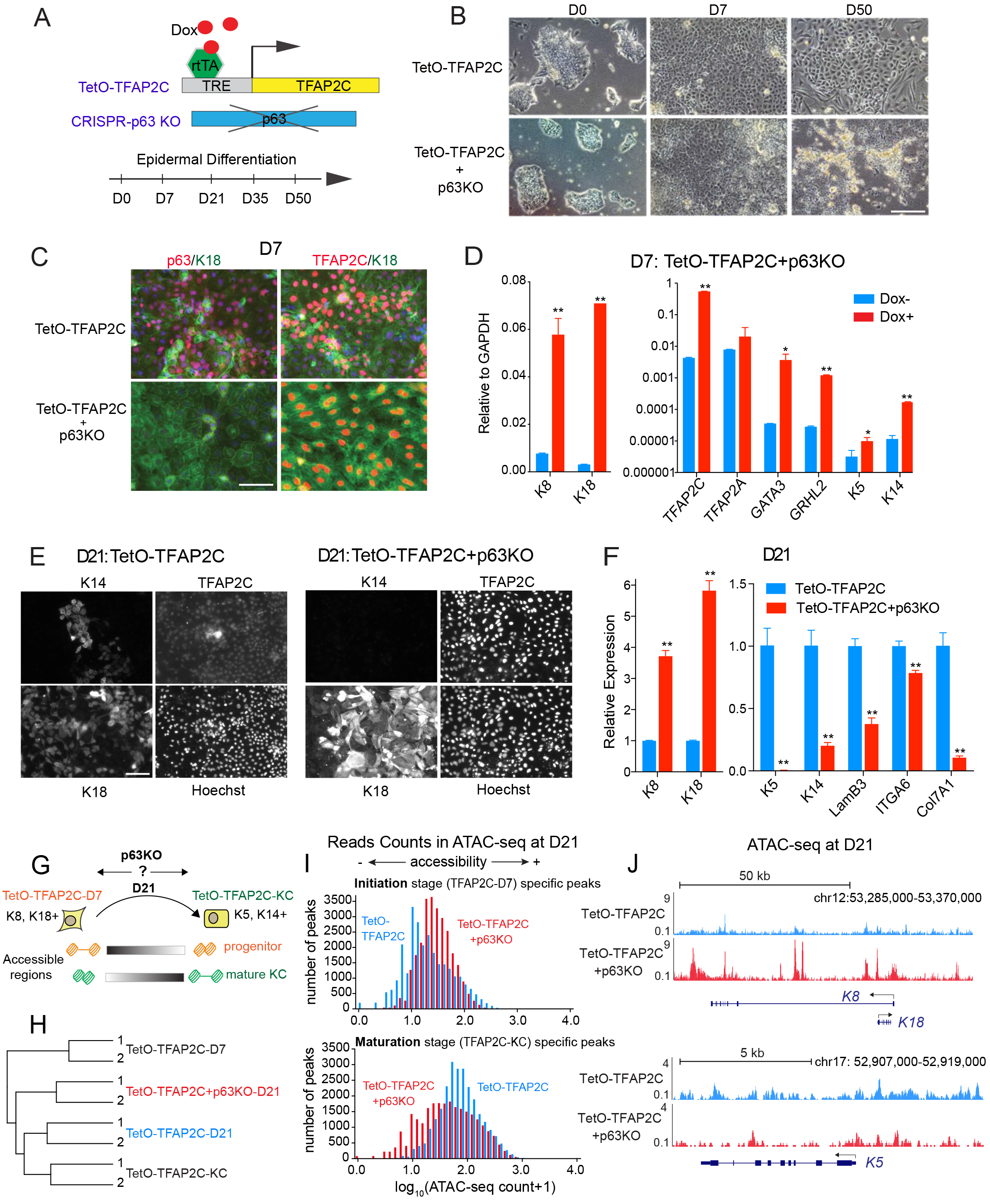
p63 Is Necessary for Keratinocyte Maturation during TFAP2C-Induced Epidermal Differentiation. (A) Schematic illustration of the approach to functionally study of p63 via CRISPR/Cas9 mediated deletion during TFAP2C-induced epidermal differentiation. (B) Morphological changes in TFAP2C-induced epidermal differentiation upon p63 deletion. Light microscopy images of the cells with genotypes labeled as “TetO-TFAP2C” and “TetO-TFAP2C+p63KO” at D0, D7 and D50 are compared, showing similar surface ectodermal epithelial morphology at D7, but strong cell death and lacking keratinocyte colonies at D50 when p63 was deleted. Scale bar: 200μm. (C-D) p63 loss of function did not affect surface ectoderm initiation at early stage of epidermal differentiation. (C) IF staining of the cells with genotypes labeled as “TetO-TFAP2C” and “TetO-TFAP2C+p63KO” at differentiation stage D7. Surface ectoderm markers K18 and TFAP2C were still highly expressed in the cells with p63 deletion. Scale bar: 50μm. (D) Gene expression analysis by qRT-PCR showing increased expression of surface ectoderm markers in TetO-TFAP2C+p63KO cells upon Dox induction. qRT-PCR values were normalized to the values of internal control GAPDH. Mean ± S.D. is shown; significant difference with *p<0.05 and **p<0.01 relative to control (Dox-) determined by t-test (n=3). (E-F) p63 loss of function resulted in failure of keratinocyte maturation at late stage of differentiation. (E) IF staining of the cells with genotypes labeled as “TetO-TFAP2C” (left panel) and “TetO-TFAP2C+p63KO” (right panel) at differentiation stage D21. The control cells (TetO-TFAP2C) are positive for mature keratinocyte maker (K14), and have lower level of surface ectoderm makers (K18, TFAP2C); while in p63 null cells (TetO-TFAP2C+p63KO), K14 is non-detectable, but K18 and TFAP2C are still highly expressed. Nuclei were stained by Hoechst. Scale bar: 40μm. (F) Gene expression analysis by qRT-PCR showing higher level of surface ectoderm markers (K8, K18) and lower level of mature keratinocyte markers (K5, K14, LamB3, ITGA6, Col7A1) upon p63 loss of function at D21. qRT-PCR values were normalized to the values of the control cells (TetO-TFAP2C). Mean ± S.D. is shown; significant difference with **p<0.01 relative to control determined by t-test (n=3). (G) Schematic illustration of the analysis of the chromatin changes upon p63 loss of function at D21. Here, we took the TetO-TFAP2C-D7(K8, K18+) and TetO-TFAP2C-KC (K5, K14+) specific accessible regions as the representative features of “progenitor” and “mature KC” respectively, and use them to evaluate the differentiation status of the cells at D21 with or without p63 (i.e. TetO-TFAP2C vs. TetO-TFAP2C+p63KO). (H) Hierarchical clustering of the ATAC-seq signals showing a close relationship between TetO-TFAP2C-D21 and mature KC (TetO-TFAP2C-KC); while loss of p63 results in cells stuck at a more immature stage between TetO-TFAP2C-D7 and TetO-TFAP2C-KC. (I) Loss of p63 resulted in higher level of chromatin accessibility at Initiation-stage specific peaks, and lower level of chromatin accessibility at Maturation-stage specific peaks. Histogram showing the distribution of ATAC-seq read counts from Initiation-stage (TetO-TFAP2C-D7, top panel) specific and Maturation-stage (TetO-TFAP2C-KC) specific peak regions in two types of cells at D21 (TetO-TFAP2C vs. TetO-TFAP2C+p63KO). (J) Genome browser tracks comparing ATAC-seq signal between TetO-TFAP2C and TetO-TFAP2C+p63KO at D21, showing a higher level of accessibility at *K8-K18* locus and a lower level at *K5* locus upon p63 loss of function. See also **Figure S5.**

To analyze p63-dependent gene expression and chromatin accessibility changes during the transition to the Maturation network, we collected D21 cells, the time point at which we previously showed were competent to induce keratinocytes (Figure 4), and checked gene expression and chromatin accessibility changes in TetO-TFAP2C+p63KO vs. TetO-TFAP2C (Figure 5G). Hierarchical clustering shows that D21 TetO-TFAP2C (i.e. TetO-TFAP2C-D21) but not p63 mutant TetO-TFAP2C cells (i.e. TetO-TFAP2C+p63KO-D21) cluster with mature KC (TetO-TFAP2C-KC), while p63 KO cells cluster with cells from time points located between TetO-TFAP2C-D7 and TetO-TFAP2C-KC. This supports an arrest of the TFAP2C-driven differentiation process in the absence of p63 (Figure 5H). Next, we took stage-specific accessible regulatory elements from TetO-TFAP2C-D7 and TetO-TFAP2C-KC as the features for epidermal progenitor and mature KC, and evaluated their openness level in differentiated cells at D21. Consistent with arrested differentiation, greater read counts appeared in progenitor-associated, rather than keratinocyte-associated, elements in p63 KO cells (TetO-TFAP2C+p63KO) compared to control cells (TetO-TFAP2C) (Figure 5I). In addition, we found significantly more overlap with progenitor elements than mature KC elements in the peak regions highly accessible in TetO-TFAP2C+p63KO than TetO-TFAP2C (Figure S5). To confirm such findings, we examined the local chromatin status at the loci of *K8-K18* and *K5*, representative genes for epidermal progenitor and mature KC. At D21, there are higher ATAC-seq signals around *K8-K18* locus, and fewer signals around *K5* locus in TetO-TFAP2C+p63KO comparing to TetO-TFAP2C (Figure 5J). Therefore, we conclude that loss of p63 arrests the TFAP2C-driven transition from progenitor program to mature KC program.

### Feedback Regulation between p63 and TFAP2C Drives the Epigenetic Transition

Our inference model indicates that keratinocytes represent a product of the transition from an TFAP2-initiated landscape to a p63-matured landscape where p63 antagonizes select TFAP2-induced changes. TFAP2C levels initially increase with concomitant increases in p63, but then drop precipitously during Maturation as p63 expression levels become TFAP2-independent (Figure 6A). To understand the mechanism of this transition, we first studied the transcriptional regulation of TFAP2C on the well-studied p63 locus (encoded by *TP63*). Previous murine and human ectodermal dysplasia patient studies demonstrate a critical regulatory element that mediates p63 positive autoregulation, known as the C40 enhancer (Figure 6B) (Antonini et al., 2006; Antonini et al., 2015). p63 binding to this key developmental enhancer mediates the dramatic increases seen during maturation (Figure 6A). Intriguingly we found the entire p63 locus inaccessible in ES and D7 TetO-TFAP2C inducible cells in the absence of Dox. By contrast, addition of Dox at D7 opened several sites within the p63 locus shown to be bound by p63, including the C40 enhancer (Figure 6B). In addition, TFAP2C ChIP-seq in TetO-TFAP2C D7 cells revealed several binding sites on the p63 locus (Figure 6B), further confirming the physical interaction of transcription factor TFAP2C with the p63 gene locus. Consistent with the findings at the p63 locus, we observe marked increase in the ATAC-seq read counts above p63 binding sites with TFAP2C expression (Figure 6C). To interrogate TFAP2C action of p63 binding sites genome wide, we examined the chromatin status of p63 binding sites previously identified in normal human keratinocytes (Zarnegar et al., 2012) and found that an overwhelming majority of the mature p63 binding sites that are closed in TetO-TFAP2C Dox-become more accessible when TFAP2C is overexpressed at D7 (Figure 6C). Taken together, we conclude that TFAP2C turns on expression of endogenous p63 during the initiation stage, and increases chromatin accessibility surrounding p63 binding sites that allows p63-dependent epidermal lineage maturation. By contrast, *TFAP2C* locus and its binding site accessibility declined during maturation (Figures S6A-B), suggesting a reciprocal regulation between the two factors.

If p63 controls the Maturation transcription factor network, then loss of p63 at D21 should result in the reduction in the accessibility of the other Maturation-associated TF and a persistence of the Initiation-associated TF binding sites. We explored TF motifs associated with differential accessibility regions upon p63 KO at D21 (Figures 6D-E). We discovered 17,057 differential peaks (accounts for 37% of total peaks found) by comparing TetO-TFAP2C+p63KO vs. TetO-TFAP2C (2-fold change, p-value < 0.01); from which 19.3% of them are peak regions gaining accessibility and 17.7% are peak regions losing accessibility (Figure 6E). Subsequent analysis revealed that motifs for Initiation TFs (AP-2gamma, AP-2alpha, GATA3 and TEAD) were enriched in higher accessible regions, while motifs for CTCF, BORIS, p63, p53 and RFX are enriched in lower accessible regions (Figure 6E) upon p63 deletion, confirming our prediction. As further confirmation, we performed PIQ analysis on TFAP2C and p63 as the representative TFs in each group and found a statistically significant increase in the binding of TFAP2C, but a decrease in the binding of p63 in p63 KO cells compared to the control cells (Figure 6F), confirming the reciprocal feedback regulation between TFAP2C and p63 during late stage of differentiation (Figure 6G).

**Figure 6.**
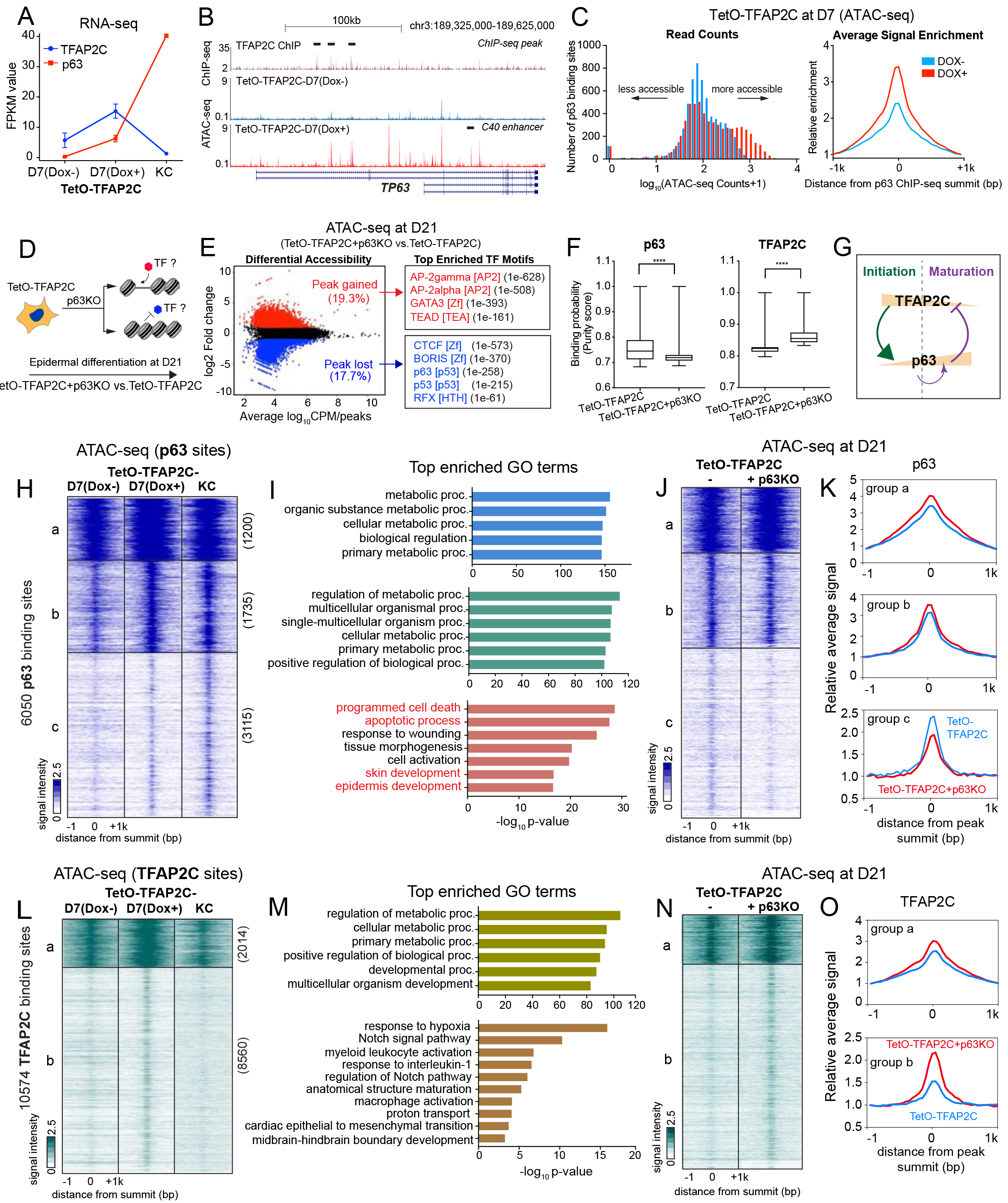
Feedback Regulation between p63 and TFAP2C Drives Chromatin Transition from Progenitor to Mature keratinocyte. (A) Gene expression changes of TFAP2C and p63 during TFAP2C-induced epidermal differentiation. The FPKM value of each gene from RNA-seq in D7 cells at progenitor stage, and “AP2C-KC” mature keratinocytes are shown. (B) Genome browser tracks comparing ATAC-seq signal between TetO-TFAP2C cells with and without Dox induction (“TetO-TFAP2C-D7 Dox+” vs. “TetO-TFAP2C-D7 Dox-”), relative to TFAP2C ChIP-seq (with black bar highlighting the peak regions) at the *TP63* locus from D7 Dox-induced differentiation. Note: two isoform types (full length and deltaN-p63) are schematically shown. There are multiple regions gaining accessibility signal in *TP63* locus upon Dox induction, including the p63 self-activation enhancer: *C40 enhancer* (black bar annotated). (C) p63 binding sites become more accessible in TFAP2C-induced surface ectoderm progenitor cells. Left panel, the histogram showing the distribution of ATAC-seq read counts within p63 ChIP-seq peak regions (Zarnegar et al., 2012) in cells with or without TFAP2C induction (“TetO-TFAP2C-D7 Dox+” vs. “TetO-TFAP2C-D7 Dox-”). Right panel, average enrichment of ATAC-seq chromatin accessibility signal within −1/+1kb region from p63 ChIP-seq peak summits in cells with or without TFAP2C induction. (D) Schematic illustration of identification of TF motifs associated with chromatin accessibility changes upon p63 loss of function. (E) Left panel: Scatter plot of differential accessibility between TetO-TFAP2C+p63KO and TetO-TFAP2C at D21. y-axis: log_2_ fold change in reads per accessible region (TetO-TFAP2C+p63KO vs. TetO-TFAP2C); x-axis: average log_10_ CPM per peak region. The values of log_2_ fold change >1, or <-1 are labeled as “peak gained” or “peak lost” with respective percentage and color. Right Panel: Top enriched TF motifs identified from differential accessible regions. TF motif name, TF family and p-values are presented. (F) PIQ footprinting analysis indicates a lower likelihood of TF occupancy of p63 and a higher likelihood of TF occupancy of TFAP2C in p63 KO cells (TetO-TFAP2C+p63KO). Significant difference with **** p<0.0001 relative to control (TetO-TFAP2C) determined by t-test. (G) A model diagram showing the negative feedback regulation between TFAP2C and p63, and the self-activation of p63 during epidermal lineage commitment. (H) Heatmap shows chromatin accessibility changes within p63-bound regions comparing samples in TFAP2C-induced epidermal differentiation. K-means clustering identified three groups of chromatin changes (labeled as “a, b, c” with binding site numbers). (I) Top enriched GO terms identified from the three clusters shown in (H) (J) Comparing the signal changes between TetO-TFAP2C+p63KO vs. TetO-TFAP2C in the defined three clusters of p63 binding sites. (K) Average enrichment of the ATAC-seq signal with −1/+1kb regions from p63 binding sites in the three clusters identified from (H) (L) Heatmap shows chromatin accessibility changes within TFAP2C-bound regions comparing samples in TFAP2C-induced epidermal differentiation. K-means clustering identified two major groups of chromatin changes (labeled as “a, b” with binding site numbers). (M) Top enriched GO terms identified from the two clusters in (L) (N) Comparing the signal changes between TetO-TFAP2+p63KO vs. TetO-TFAP2C in the defined two clusters of TFAP2C binding sites. (O) Average enrichment of the ATAC-seq signal with −1/+1kb regions form TFAP2C binding sites in the two clusters identified from (L). See also **Figure S6.**

To identify which sets of genes are subjected to TFAP2C/p63 feedback regulation genome-wide, we analyzed the p63 and TFAP2C ChIP-seq data, chromatin accessibility and gene expression at TetO-TFAP2C-D7 Dox-, Dox+ and TetO-TFAP2C-KC during TFAP2C-induced keratinocyte differentiation. The inference model predicts the existence of at least three types of p63 binding sites: a: Sites unchanged by TFAP2C or p63; b: Sites opened by TFAP2C that stay open independent of p63; c: Sites opened by TFAP2C and opened further by elevated p63 protein levels. Indeed, analysis of the chromatin accessibility changes at p63 binding sites revealed the predicted three groups of regions (a, b, c) (Figures 6H-I, and S6C). Group a p63 binding site consists of metabolic genes that are constitutively open regions in all three samples, and are located in promoter-proximal regulatory regions, and Group b genes that are located in both proximal and distal regions, and depend only on early TFAP2C for accessibility. By contrast, Group c sites are keratinocyte maturation genes containing GO terms “programmed cell death”, “response to wounding”, “skin development”, “epidermis development”. While they demonstrate a small increase in accessibility with TFAP2C expression, there is a strong dependence on high level p63 expression to achieve the marked accessibility seen in mature keratinocytes. These findings indicate that key keratinocyte maturation genes are “primed” early by TFAP2C and then become fully accessible through increased p63 expression during maturation. A test of this prediction is that in TetO-TFAP2C+p63KO cells at D21, Group c sites should fail to increase or decrease as TFAP2C levels fall and p63 is null. Indeed, while Group a and b genes show no change in accessibility in the absence of p63, Group c genes demonstrate decreased accessibility in the absence of p63 (Figures 6J-K).

Reciprocal regulation by p63 of TFAP2C D7 binding site accessibility illuminates p63 feedback regulation on a subset of the TFAP2C network. We identified two groups of TFAP2C binding sites (Figures 6L-M and S6D): Group a TFAP2C sites consist of metabolic genes that are located in both proximal and distal regulatory regions, while Group b sites consist of genes with GO terms “response to hypoxia”, “Notch signaling pathway” and “epithelial to mesenchymal transition”, and are located in distal regions. In both groups, binding site accessibility increases with TFAP2C expression and decreases with decreasing TFAP2C levels in late differentiation (Figure 6L). Consistent with p63 negative feedback regulation of TFAP2C, the accessible signal of both groups significantly increases upon p63 deletion, with group b demonstrating much greater sensitivity to p63 levels (Figures 6N-O). We conclude that p63 increases accessibility of key epidermal genes as its levels increase, while shutting down the accessibility of a subset of TFAP2C binding sites associated with simple epithelium.

## DISCUSSION

In this study, we use paired temporal chromatin accessibility and transcriptome data from hESC-derived epidermal differentiation to identify two distinct TF-chromatin regulatory networks responsible for epidermal lineage initiation and maturation. Through functional validation we make the surprising finding that TFAP2C acts as the “Initiation factor” which induces the surface ectoderm chromatin landscape on which p63 subsequently acts as the “maturation factor” to modify the pre-patterned landscape into that of functional keratinocytes. TFAP2C induces p63 expression and primes p63 binding site accessibility, allowing p63 positive autoregulation and negative feedback regulation to close select TFAP2C binding sites, therefore driving landscape maturation. The two-step, feed-forward autoregulation and negative feedback regulatory mechanisms between Initiation and Maturation factors provide a novel and general logic to understand the chromatin dynamics of tissue lineage commitment (Figure 7).

**Figure 7.**
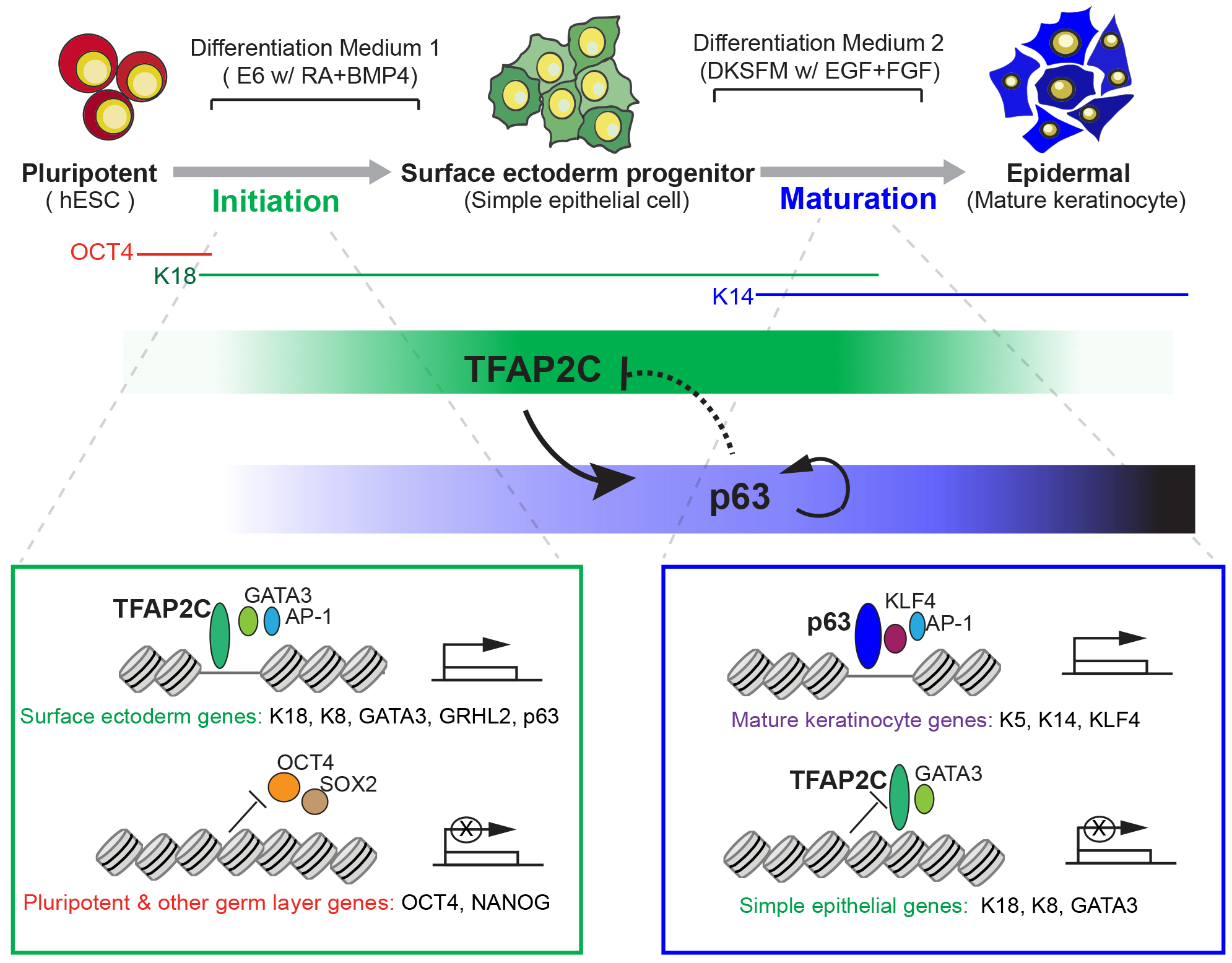
A Model of Epigenomic Regulation during Epidermal Lineage Commitment. A proposed model depicts the identified chromatin states and feedback regulation between TFAP2C- and p63-centered TF regulatory network driving the chromatin transition during epidermal lineage commitment.

Our network model provides several advantages to understand lineage commitment at the genome-wide level. First, it integrates matched chromatin accessibility, transcription factor binding, gene expression, and histone modifications into a “TF-open chromatin regulatory element-TG” triplet, greatly facilitating accurate inference of gene regulatory relations. The paired gene expression and chromatin accessibility data help filter out noise inherent in motif site analysis, and improves on the existing “footprinting” strategy that depends on the shape of the DNase-seq (or ATAC-seq) profile. In addition, our approach incorporates publicly available chromatin conformation data that allows accurate assignment of dynamically changing distal enhancer accessible sites to their TGs. In the current iteration, we used several data sources including distance of the RE to the TG, chromosomal conformation data, such as Hi-C, that reveals the connection of distal DNA element to promoter, and the co-occurrence of chromatin openness between each distal element and promoter pairs from ENCODE DHS datasets in variety of tissues. However, we anticipate the incorporation of newer conformation technologies, such as ChIA-PET and Hi-CHIP (Fullwood et al., 2009; Mumbach et al., 2016) will improve the coverage of RE and TG association and deepen the inferred TF network.

Gain and loss of function studies reveal the surprising finding that the early and late networks can be collapsed into the actions of TFAP2C and p63. Previous studies have revealed important functions of TFAP2C in various biological process, such as trophectodermal development (Auman et al., 2002), and breast cancer progress (Woodfield et al., 2010). In skin development, loss of Tfap2c results in disrupted epidermal gene expression and delayed skin development (Guttormsen et al., 2008; Qiao et al., 2012), demonstrating an essential role in epidermal lineage commitment. Our ATAC-seq and ChIP-seq data support the hypothesis that TFAP2C acts directly on chromatin to alter chromatin accessibility. This data is consistent with previous reports of TFAP2 family members possessing pioneer factor characteristics (Pihlajamaa et al., 2014; Tan et al., 2011).

By contrast, TFAP2C in other systems requires chromatin regulators that act as co-activators or co-repressors in TFAP2C-mediated gene regulation (Braganca et al., 2003; Braganca et al., 2002; Wong et al., 2012), leading us to speculate that TFAP2C might recruit specific chromatin regulators to achieve surface ectoderm gene activation. TFAP2C binding preferentially to active chromatin regions marked by H3K27ac and H3K4m1 distal to the TSS suggests it acts through long range enhancer-promoter interaction and enhancer-associated chromatin factors, such as p300/CBP, MLL/Set1 family and chromatin remodeler (Calo and Wysocka, 2013). While our work shows the necessary and sufficiency for TFAP2C to induce the Initiation TF network, potential cross regulatory interactions with TFAP2C targets like GATA3 or GRHL2 prevent us from excluding a role for the network TFs in modulating or synergizing with TFAP2C to affect the ultimate composition of the dynamic chromatin landscape. This includes a role for other TFAP2 family members, which share cross-regulation of each other’s loci and can homodimerize with each other, complicating a straightforward genetic epistasis analysis.

Besides inducing surface ectoderm commitment, a major insight of this work is the demonstration that TFAP2C prepares the underlying chromatin landscape for p63-dependent keratinocyte maturation. TFAP2C activates the transcription of, and increases the binding site accessibility for, p63. p63, in turn, modifies the TFAP2C-mediated chromatin landscape toward epidermal maturation through self-activation. Our work tempers the long-held view that p63 is the master regulator of keratinocyte differentiation, and places p63 as a critical but dependent partner to the TFAP2C-modified landscape. p63 plays a central role in the development of a wide variety of ectodermal and endodermal-derived epithelia, including epidermal development (Mills et al., 1999; Yang et al., 1999), mammary, lung, and prostate (Crum and McKeon, 2010), but remarkably, loss of p63 in TFAP2C-induced surface ectoderm differentiation or forced expression in hPSCs (Medawar et al., 2008) does not affect differentiation. Our work shows that the genome context on which p63 acts ultimately determines the final differentiation product, and suggests that small changes in diffusible morphogens RA/BMP could dramatically alter TFAP2C levels, the set of enhancer sites that are p63-sensitive (group c, Figure 6H), and the gene expression profile of the epidermis generated. Our results also explain how a single lineage selector like p63 can direct such a panoply of transcriptional programs during development, as it is expressed in the context of distinct pre-established chromatin landscapes.

A surprising revelation from the network modeling was that p63 drives maturation both by positive autoregulation of its own locus and by inhibition of the immature surface epithelial landscape. In response to maturation medium, p63 levels are dramatically boosted by self-activation from the p63 C40 enhancer locus (Antonini et al., 2006; Antonini et al., 2015), facilitating the activation of keratinocyte specific targets. By contrast p63 reduces the level and binding site accessibility of TFAP2C in at least two ways. P63 represses the transcription of TFAP2C itself. We have found a p63 binding site located distal to the TSS of TFAP2C and p63 inhibits the expression through looping with TSS locus (Figure S6A, and unpublished data). Also, p63 antagonizes a subset of TFAP2C TG activated during initiation (Figures 6L-O). Interestingly, TFAP2C remains required at particular loci for mature keratinocyte function (McDade et al., 2012), arguing that p63 action must repress only a subset of potential targets and that the balance between TFAP2C and p63 networks determines the nature of the mature end product.

Taken together, our work provides a general analysis strategy to depict epigenetic transition during lineage commitment through TF-chromatin network modeling, and reveals a new regulatory principle between lineage initiation factor and maturation factors to ensure faithful tissue generation. Such findings will bring more insights to understand the logic of somatic tissue development, and improve the assessment of stem cell competency and differentiation efficiency in tissue engineering and regenerative medicine.

## EXPERIMENTAL PROCEDURES

### Cell culture and differentiation

H9 human ESCs and modified cell lines were seeded in a feeder-free system using Matrigel hESC-Qualified Matrix (BD Corning) and were maintained in Essential 8 media (Life Technologies). To induce differentiation into keratinocyte, the cells were first fed with “differentiation medium 1” containing 5ng/ml BMP4 and 1 μM RA in Essential 6 media (Life Technologies) for 7 days; thus forming surface ectodermal progenitor cells. After seven days, the medium was changed to “differentiation medium 2”, i.e. Defined Keratinocyte Serum Free Medium (DKSFM) with growth supplements containing insulin, EGF and FGF (Life Technologies), and the cells underwent selection and expansion for 2 months to get maturation into keratinocyte. The resulting keratinocyte colonies were passaged onto Corning PureCoat™ ECM Mimetic 6-well Collagen I Peptide Plate (Corning) and expanded in DKSFM medium. See **Supplemental Experimental Procedure** for more details.

### In vitro skin reconstitution assay

Generation of organotypic epidermis was performed by following the protocol described previously (Sebastiano et al., 2014) with minor modification. Basically, 1X10^6^ keratinocytes derived from normal differentiation with H9 hESC or from TFAP2C-induced differentiation were collected and seeded on top of devitalized dermis and cultured submerged in DKSF medium for 5 days. The medium was then gradually changed to Keratinocyte Growth Medium for 7 days, after which stratification was induced by raising the dermis to air-liquid interface. After 2 weeks, the reconstituted epidermis was collected for IF staining.

### RNA-seq, ChIP-seq and ATAC-seq

RNA extraction was performed using RNeasy Plus (Qiagen) from the samples reported in this research. Ribosomal RNA was removed from each RNA extraction using Ribo-Zero Gold rRNA Removal kit (Illumina). The RNA-seq libraries were constructed by TruSeq Stranded mRNA Library Prep kit (Illumina). ChIP assay was performed following previously described method (Calo et al., 2015) with minor modifications. The libraries were prepared following the NEBNext protocol. ATAC-seq was performed as described (Buenrostro et al., 2013) All the libraries were sequenced to saturation on Illumina Hiseq2000 or NextSeq sequencers. More details of the sequencing library preparation and data processing can be found in **Supplemental Experimental Procedure.**

### TF-RE-TG Triplet Inference Modeling

We developed a statistical model to integrate ATAC-seq and RNA-seq data and to infer, from the observed expression and accessibility data in time-course cellular context, how each RE interacts with relevant TFs to affect the expression of its TGs. Our strategy was to identify the upstream TFs and downstream genes for a RE. We next assembled a union file of peaks called from ATAC-seq across all time samples. Next, we treated each TF-RE-TG triplet as the basic regulatory unit (Figure 2B)., ranked them by integrating genomic features and extracted significant regulatory relations by maximizing the joint probability P(TF, RE, TG)=P(TG|RE)P(RE|TF)P(TF). Several assumptions were made. RE openness was defined as the fold enrichment of the read starts in this region versus the read starts in a 1Mb background window. Next, each TF was described by its motif binding score to the RE and its expression value, which was the FPKM value from RNA-seq experiments. The regulation of a TF on an RE was quantified as the regression of TF expression to RE openness. Finally, regulation from RE to TG used sources that correlate the openness and TG expression (ENCODE) plus some constraints to identify direct regulation. In the time course differentiation dataset, we observed three major stages by the global dynamics of chromatin state. Stage 1 has one sample (D0), stage 2 has three samples (D7, D14, D21) and stage 3 has two samples (D43 and H9KC). Therefore, we reconstructed the TF-RE-TG network to integrate dynamic change of driver TFs and downstream TG with union peak elements. We proposed a model to infer the RE’s openness change from Stage 1 to 2 (surface ectoderm initiation) and Stage 2 to 3 (keratinocyte maturation). The expression level for each stage was taken as the maximum when there were multiple samples. Our computation has four major steps: 1. Predicting RE’s target gene; 2. Collecting genomic features from chromatin state and expression level; 3. Integrating genomic features and rank TF-RE-TG triplets; 4. Controlling FDR and pooling significant TF-RE-TG triplets into network. The networks are visualized by Cytoscape (Shannon et al., 2003). Each transition period contains two regulatory networks representing the TF regulation changes on open and closed chromatin regions associated with a particular TG. The full list of TF-RE-TG triplets from each network is shown in Table S3. More detailed analysis pipeline can be found in **Supplemental Experimental Procedure.**

## DATA AND SOFTWARE AVAILABILITY

The deep sequencing dataset (including ATAC-seq, RNA-seq and ChIP-seq) in this study, are deposited in Gene Expression Omnibus database (accession number GEO: GSE108248).

## SUPPLEMENTAL INFORMATION

Supplemental Information includes six **Supplemental Figures**, three **Supplemental Tables** and **Supplemental Experimental Procedure**.

## AUTHOR CONTRIBUTIONS

Conceptualization, Methodology, Writing, L.L., Y.W., W.H.W., and A.E.O.; Investigation, L.L., J.L.T., J.M.P., H.H.Z., F.F., S.P.M., S.N.P., and R.L.; Software and Formal analysis, Y.W., G.S., Z.D., J.L., E.J.L., and L.C.; Data Curation, G.S., and Y.W.; Resource, M.W., H.Y.C., W.H.W., and A.E.O.; Funding Acquisition, H.Y.C., W.H.W., and A.E.O.; Supervision, H.Y.C., W.H.W., and A.E.O.

## ACKNOWLEDGEMENTS

We thank Stanford Functional Genomics Facility for support with deep sequencing study; S. Yamanaka for sharing PiggyBac inducible expression plasmid; E. Bashkirova, M. Ameen, S. Tao and K. Qu for technical support. We thank P. Khavari, A. Christiano, E. Fuchs, and Q. Luo for pre-submission review, and other Oro and Chang laboratory members for discussion and support. We thank I. Gitman for administrative assistance. This work was supported by NIH Grants P50-HG007735 (to H.Y.C, and W.H.W), R01GM109836 (to W.H.W), and F32AR070565 (to J.M.P.), by California Institute for Regenerative Medicine Tools Grant RT3-07796 (to A.E.O), by EB Research Partnership (to A.E.O), and by National Natural Science Foundation of China Grants 61671444 and 61621003 (Y.W.)

## REFERENCES

Antonini, D., Rossi, B., Han, R., Minichiello, A., Di Palma, T., Corrado, M., Banfi, S., Zannini, M., Brissette, J.L., and Missero, C. (2006). An autoregulatory loop directs the tissue-specific expression of p63 through a long-range evolutionarily conserved enhancer. Mol Cell Biol 26, 3308–3318.

Antonini, D., Sirico, A., Aberdam, E., Ambrosio, R., Campanile, C., Fagoonee, S., Altruda, F., Aberdam, D., Brissette, J.L., and Missero, C. (2015). A composite enhancer regulates p63 gene expression in epidermal morphogenesis and in keratinocyte differentiation by multiple mechanisms. Nucleic Acids Res 43, 862–874.

Auman, H.J., Nottoli, T., Lakiza, O., Winger, Q., Donaldson, S., and Williams, T. (2002). Transcription factor AP-2gamma is essential in the extra-embryonic lineages for early postimplantation development. Development 129, 2733–2747.

Braganca, J., Eloranta, J.J., Bamforth, S.D., Ibbitt, J.C., Hurst, H.C., and Bhattacharya, S. (2003). Physical and functional interactions among AP-2 transcription factors, p300/CREB-binding protein, and CITED2. J Biol Chem 278, 16021–16029.

Braganca, J., Swingler, T., Marques, F.I., Jones, T., Eloranta, J.J., Hurst, H.C., Shioda, T., and Bhattacharya, S. (2002). Human CREB-binding protein/p300-interacting transactivator with ED-rich tail (CITED) 4, a new member of the CITED family, functions as a co-activator for transcription factor AP-2. J Biol Chem 277, 8559–8565.

Buenrostro, J.D., Giresi, P.G., Zaba, L.C., Chang, H.Y., and Greenleaf, W.J. (2013). Transposition of native chromatin for fast and sensitive epigenomic profiling of open chromatin, DNA-binding proteins and nucleosome position. Nat Methods 10, 1213–1218.

Calo, E., Flynn, R.A., Martin, L., Spitale, R.C., Chang, H.Y., and Wysocka, J. (2015). RNA helicase DDX21 coordinates transcription and ribosomal RNA processing. Nature 518, 249–253.

Calo, E., and Wysocka, J. (2013). Modification of enhancer chromatin: what, how, and why? Mol Cell 49, 825–837.

Crum, C.P., and McKeon, F.D. (2010). p63 in epithelial survival, germ cell surveillance, and neoplasia. Annu Rev Pathol 5, 349–371.

Duren, Z., Chen, X., Jiang, R., Wang, Y., and Wong, W.H. (2017). Modeling gene regulation from paired expression and chromatin accessibility data. Proc Natl Acad Sci U S A 114, E4914–E4923.

Ernst, J., and Kellis, M. (2012). ChromHMM: automating chromatin-state discovery and characterization. Nat Methods 9, 215–216.

Fullwood, M.J., Liu, M.H., Pan, Y.F., Liu, J., Xu, H., Mohamed, Y.B., Orlov, Y.L., Velkov, S., Ho, A., Mei, P.H., et al. (2009). An oestrogen-receptor-alpha-bound human chromatin interactome. Nature 462, 58–64.

Guenou, H., Nissan, X., Larcher, F., Feteira, J., Lemaitre, G., Saidani, M., Del Rio, M., Barrault, C.C., Bernard, F.X., Peschanski, M., et al. (2009). Human embryonic stem-cell derivatives for full reconstruction of the pluristratified epidermis: a preclinical study. Lancet 374, 1745–1753.

Guttormsen, J., Koster, M.I., Stevens, J.R., Roop, D.R., Williams, T., and Winger, Q.A. (2008). Disruption of epidermal specific gene expression and delayed skin development in AP-2 gamma mutant mice. Dev Biol 317, 187–195.

Koster, M.I., and Roop, D.R. (2007). Mechanisms regulating epithelial stratification. Annu Rev Cell Dev Biol 23, 93–113.

Li, L., Liu, C., Biechele, S., Zhu, Q., Song, L., Lanner, F., Jing, N., and Rossant, J. (2013). Location of transient ectodermal progenitor potential in mouse development. Development 140, 4533–4543.

McDade, S.S., Henry, A.E., Pivato, G.P., Kozarewa, I., Mitsopoulos, C., Fenwick, K., Assiotis, I., Hakas, J., Zvelebil, M., Orr, N., et al. (2012). Genome-wide analysis of p63 binding sites identifies AP-2 factors as co-regulators of epidermal differentiation. Nucleic Acids Res 40, 7190–7206.

Medawar, A., Virolle, T., Rostagno, P., de la Forest-Divonne, S., Gambaro, K., Rouleau, M., and Aberdam, D. (2008). DeltaNp63 is essential for epidermal commitment of embryonic stem cells. PLoS One 3, e3441.

Metallo, C.M., Ji, L., de Pablo, J.J., and Palecek, S.P. (2008). Retinoic acid and bone morphogenetic protein signaling synergize to efficiently direct epithelial differentiation of human embryonic stem cells. Stem Cells 26, 372–380.

Mills, A.A., Zheng, B., Wang, X.J., Vogel, H., Roop, D.R., and Bradley, A. (1999). p63 is a p53 homologue required for limb and epidermal morphogenesis. Nature 398, 708–713.

Mumbach, M.R., Rubin, A.J., Flynn, R.A., Dai, C., Khavari, P.A., Greenleaf, W.J., and Chang, H.Y. (2016). HiChIP: efficient and sensitive analysis of protein-directed genome architecture. Nat Methods 13, 919–922.

Pihlajamaa, P., Sahu, B., Lyly, L., Aittomaki, V., Hautaniemi, S., and Janne, O.A. (2014). Tissue-specific pioneer factors associate with androgen receptor cistromes and transcription programs. EMBO J 33, 312–326.

Qiao, Y., Zhu, Y., Sheng, N., Chen, J., Tao, R., Zhu, Q., Zhang, T., Qian, C., and Jing, N. (2012). AP2gamma regulates neural and epidermal development downstream of the BMP pathway at early stages of ectodermal patterning. Cell Res 22, 1546–1561.

Qu, Y., Zhou, B., Yang, W., Han, B., Yu-Rice, Y., Gao, B., Johnson, J., Svendsen, C.N., Freeman, M.R., Giuliano, A.E., et al. (2016). Transcriptome and proteome characterization of surface ectoderm cells differentiated from human iPSCs. Sci Rep 6, 32007.

Rao, S.S., Huntley, M.H., Durand, N.C., Stamenova, E.K., Bochkov, I.D., Robinson, J.T., Sanborn, A.L., Machol, I., Omer, A.D., Lander, E.S., et al. (2014). A 3D map of the human genome at kilobase resolution reveals principles of chromatin looping. Cell 159, 1665–1680.

Sahoo, D., Dill, D.L., Tibshirani, R., and Plevritis, S.K. (2007). Extracting binary signals from microarray time-course data. Nucleic Acids Res 35, 3705–3712.

Sebastiano, V., Zhen, H.H., Haddad, B., Bashkirova, E., Melo, S.P., Wang, P., Leung, T.L., Siprashvili, Z., Tichy, A., Li, J., et al. (2014). Human COL7A1-corrected induced pluripotent stem cells for the treatment of recessive dystrophic epidermolysis bullosa. Science translational medicine 6, 264ra163.

Shannon, P., Markiel, A., Ozier, O., Baliga, N.S., Wang, J.T., Ramage, D., Amin, N., Schwikowski, B., and Ideker, T. (2003). Cytoscape: a software environment for integrated models of biomolecular interaction networks. Genome Res 13, 2498–2504.

Sherwood, R.I., Hashimoto, T., O’Donnell, C.W., Lewis, S., Barkal, A.A., van Hoff, J.P., Karun, V., Jaakkola, T., and Gifford, D.K. (2014). Discovery of directional and nondirectional pioneer transcription factors by modeling DNase profile magnitude and shape. Nat Biotechnol 32, 171–178.

Tan, S.K., Lin, Z.H., Chang, C.W., Varang, V., Chng, K.R., Pan, Y.F., Yong, E.L., Sung, W.K., and Cheung, E. (2011). AP-2gamma regulates oestrogen receptor-mediated long-range chromatin interaction and gene transcription. EMBO J 30, 2569–2581.

Thurman, R.E., Rynes, E., Humbert, R., Vierstra, J., Maurano, M.T., Haugen, E., Sheffield, N.C., Stergachis, A.B., Wang, H., Vernot, B., et al. (2012). The accessible chromatin landscape of the human genome. Nature 489, 75–82.

Umegaki-Arao, N., Pasmooij, A.M., Itoh, M., Cerise, J.E., Guo, Z., Levy, B., Gostynski, A., Rothman, L.R., Jonkman, M.F., and Christiano, A.M. (2014). Induced pluripotent stem cells from human revertant keratinocytes for the treatment of epidermolysis bullosa. Science translational medicine 6, 264ra164.

Wenzel, D., Bayerl, J., Nystrom, A., Bruckner-Tuderman, L., Meixner, A., and Penninger, J.M. (2014). Genetically corrected iPSCs as cell therapy for recessive dystrophic epidermolysis bullosa. Science translational medicine 6, 264ra165.

Wong, P.P., Miranda, F., Chan, K.V., Berlato, C., Hurst, H.C., and Scibetta, A.G. (2012). Histone demethylase KDM5B collaborates with TFAP2C and Myc to repress the cell cycle inhibitor p21(cip) (CDKN1A). Mol Cell Biol 32, 1633–1644.

Woodfield, G.W., Chen, Y., Bair, T.B., Domann, F.E., and Weigel, R.J. (2010). Identification of primary gene targets of TFAP2C in hormone responsive breast carcinoma cells. Genes Chromosomes Cancer 49, 948–962.

Yang, A., Schweitzer, R., Sun, D., Kaghad, M., Walker, N., Bronson, R.T., Tabin, C., Sharpe, A., Caput, D., Crum, C., et al. (1999). p63 is essential for regenerative proliferation in limb, craniofacial and epithelial development. Nature 398, 714–718.

Zarnegar, B.J., Webster, D.E., Lopez-Pajares, V., Vander Stoep Hunt, B., Qu, K., Yan, K.J., Berk, D.R., Sen, G.L., and Khavari, P.A. (2012). Genomic profiling of a human organotypic model of AEC syndrome reveals ZNF750 as an essential downstream target of mutant TP63. Am J Hum Genet 91, 435–443.

